# Heterogeneity in Nucleosome Spacing Governs Chromatin Elasticity

**DOI:** 10.1101/708966

**Authors:** Bruno Beltran, Deepti Kannan, Quinn MacPherson, Andrew J. Spakowitz

## Abstract

Within a living cell, the myriad of proteins that bind DNA introduce heterogeneously spaced kinks into an otherwise semiflexible DNA double helix. To investigate the effects of heterogeneous nucleosome binding on chromatin organization, we extend the wormlike chain (WLC) model to include statistically spaced, rigid kinks. On time scales where nucleosome positions are fixed, we find that the probability of chromatin loop formation can differ by up to six orders of magnitude between two sets of nucleosome positions drawn from the same distribution. On longer time scales, we show that continuous re-randomization due to nucleosome turnover results in chromatin tracing out an effective WLC with a dramatically smaller Kuhn length than bare DNA. Together, these observations demonstrate that heterogeneity in nucleosome spacing acts as the dominant source of chromatin elasticity and governs both local and global chromatin organization.

---

The spatial organization of chromatin—genomic DNA and its associated proteins—is critical to a range of biological processes, from controlling gene expression [1] to facilitating DNA damage repair [2, 3]. The fundamental unit of eukaryotic chromatin organization is the nucleosome, which consists of 147 basepairs of DNA wrapped around a histone-protein octamer [4]. Linker DNA connecting adjacent nucleosomes ranges from less than 10 bp on average in fission yeast [5] to more than 80 bp in sea urchin sperm cells [6].

*In vitro* images of chromatin have historically contained regular, “30-nm fiber” helical structures, motivating models of chromatin with constant (periodic) nucleosome spacing [7–16]. However, recent super-resolution microscopy measurements indicate that *in vivo*, interphase mammalian chromatin instead forms a disordered, “beads-on-a-string” structure [17–19]. In addition, the latest *in vivo* nucleosome positioning data suggest that linker lengths are extremely heterogeneous. The occupancy profiles of even the most well-positioned nucleosomes—such as those near transcription start sites—are well described by a model where nucleosomes bind nearly uniformly along the DNA [11, 20–30] and are merely excluded from certain areas [31].

So far, works that address linker length heterogeneity have been either simulation studies [32, 33] or purely geometrical models [34, 35]. These models can produce individual “beads-on-a-string” configurations qualitatively similar to those observed in bulk chromatin. However, there remains a need for an analytical approach that can systematically characterize how and when linker length heterogeneity leads to structural disorder.

In this paper, we characterize the structures that emerge from the combined effects of thermal fluctuations in the DNA linkers and the geometric effects of experimentally-relevant linker length heterogeneity. We show that almost any linker length heterogeneity is sufficient to produce the types of disordered chromatin structures that are now believed to dominate nuclear architecture. The intuition behind our structural claims extends to any polymer composed of aperiodic kinks, such as the dihedral “kinks” found at junctions of a block copolymer. More broadly, our results contribute to a large class of problems in which quenched disorder competes with thermal fluctuations to determine the structural properties of a system.

We model each DNA linker as a twistable wormlike chain (TWLC), and the nucleosomes as the points where these linker strands connect. At each nucleosome, we impose the constraint that the incoming and outgoing linker DNA strands have a fixed relative orientation. The orientation of the strand entering the *i*th nucleosome is defined by the matrix 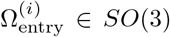 that rotates the lab frame into the entry orientation. The exit orientation 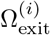 is related to the entry orientation by a kink rotation Ω_kink_, such that 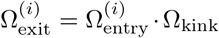, as shown in Fig. 1.

**FIG. 1.**
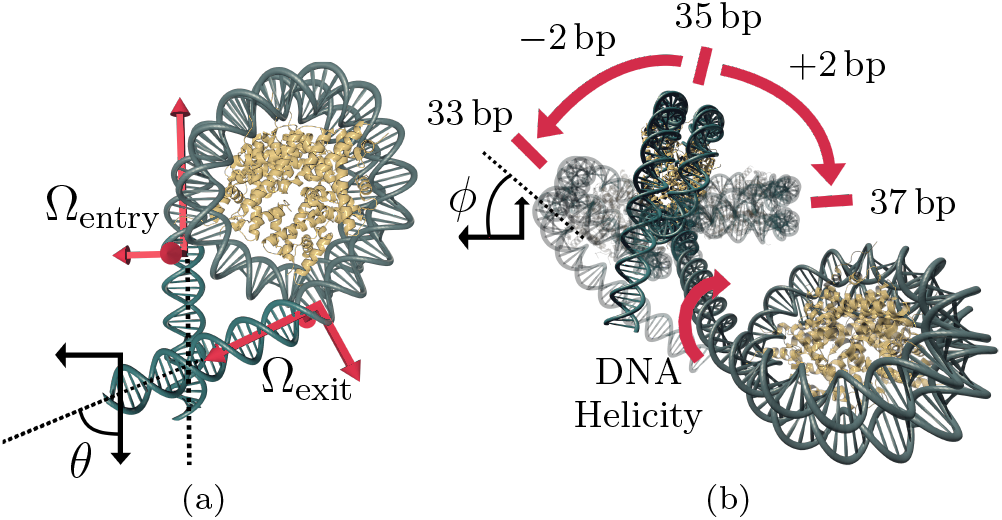
(a) The structure of a human nucleosome [43] with straight linkers extrapolated from the entry (Ωentry) and exit (Ωexit) orientations of the bound DNA. The amount of DNA wrapping the nucleosome dictates the spherical angle *θ*. (b) Two adjacent nucleosomes at zero temperature. The DNA double helix has an intrinsic twist density (*τ* = 2*π/*(10.5 bp)). If we anchor the location of one nucleosome, the binding orientation of the next histone octamer must change so that it aligns with the major groove of the double helix. This means that as the linker length *L* connecting two nucleosomes gets longer or shorter, the relative orientations of adjacent octamers changes to create an angle *φ* = *τ L*.

We represent a TWLC of length *L* as a space curve 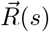, *s* ∈ [0, *L*]. The chain orientation at each point along this curve, Ω(*s*), is represented by an orthonormal triad 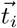, where 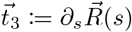. We track the bend and twist of our polymer via the angular “velocity” vector 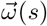, which operates as 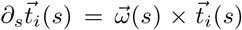. The Green’s function of the first linker represents the probability that a polymer of length *L*_1_ that begins at the origin with fixed initial orientation Ω_0_ ends at position 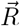 with fixed end orientation Ω. For a TWLC with no kinks, the Green’s function is given by

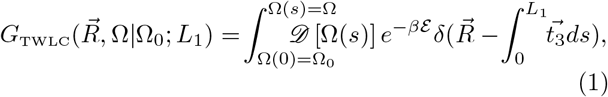

where the energy

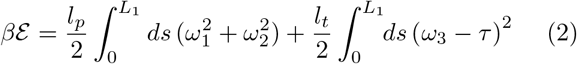

is quadratic in bending and twisting deformation. The natural twist of DNA gives *τ* = 2*π*(10.5 bp)^−1^, and we set the persistence length *l*_*p*_ = 50 nm and twist persistence length *l*_*t*_ = 100 nm to match measurements of DNA elasticity [36–39].

Reference [40] solves Equation 1 analytically in Fourier space 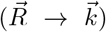 by computing the coefficients in the Wigner D-function expansion

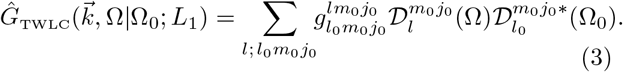

To account for the kink introduced by the nucleosome, we rotate the final orientation of the linker DNA, Ω = Ω_entry_, to Ω_exit_ using the formula

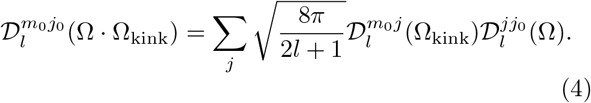

The resulting Green’s function combines the effects of a linker DNA and a nucleosome, but is still a Wigner Dfunction expansion with modified coefficients 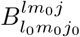 (first computed in Ref. [41]). We present an alternative derivation in Supplemental Material [42].

X-ray crystallography [4, 44, 45] and cryo-EM [43, 46–49] measurements of the nucleosome show that histone-bound DNA is well approximated by a deformed B-DNA structure, wrapping the histone octamer 1.7 times in a superhelix with a radius of 4.19 nm and a pitch of 2.59 nm [45]. Thus, Ω_entry_ and Ω_exit_ are well defined as a function of the number of bound nucleotides to the histone core. In what follows, we fix the wrapping level to that found in the crystal structure (147 bp). Using different values for the wrapping level rescales our results (see Supplemental Material [42], Fig. S3 and Fig. S10).

To compose monomers of the nucleosome chain with prescribed linker lengths, we perform an iterated convolution of the Green’s function for each nucleosome-linker pair. In Fourier space, this corresponds to multiplying the matrices 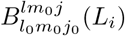.

A key property of our model is that the relative orientation of adjacent nucleosomes is not only determined by Ω_kink_ and the thermal fluctuations of the linker strand, but also by changing the length of the linker strand (as demonstrated in Fig. 1b). Our propagator *G* takes this into account implicitly due to the inclusion of *τ* in Eq. 2.

We begin by computing the end-to-end distance 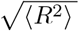 of the chain using the formula 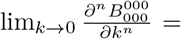 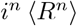. From this, we extract the Kuhn length, 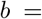 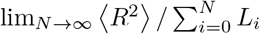, which gives a universal measure of the elasticity of a polymer at long length scales.

In Fig. 2a, we plot 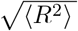 as a function of chain length for homogeneous chains of nucleosomes with 36 bp and 38 bp linkers. We compare each of these curves to the 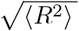 of a TWLC with the same Kuhn length but without kinks. At short length scales, the initial slope of 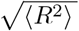 for all chains is one, corresponding to rigid-rod behavior. At chain lengths comparable to the persistence length, the bare WLC’s slope smoothly transitions to 1/2 (on a log-log scale), corresponding to random-walk behavior. In contrast, the homogeneous chain 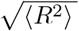 jumps from that of bare DNA to that of the best-fit WLC, whose Kuhn length is dramatically smaller than twice the persistence length of bare DNA.

**FIG. 2.**
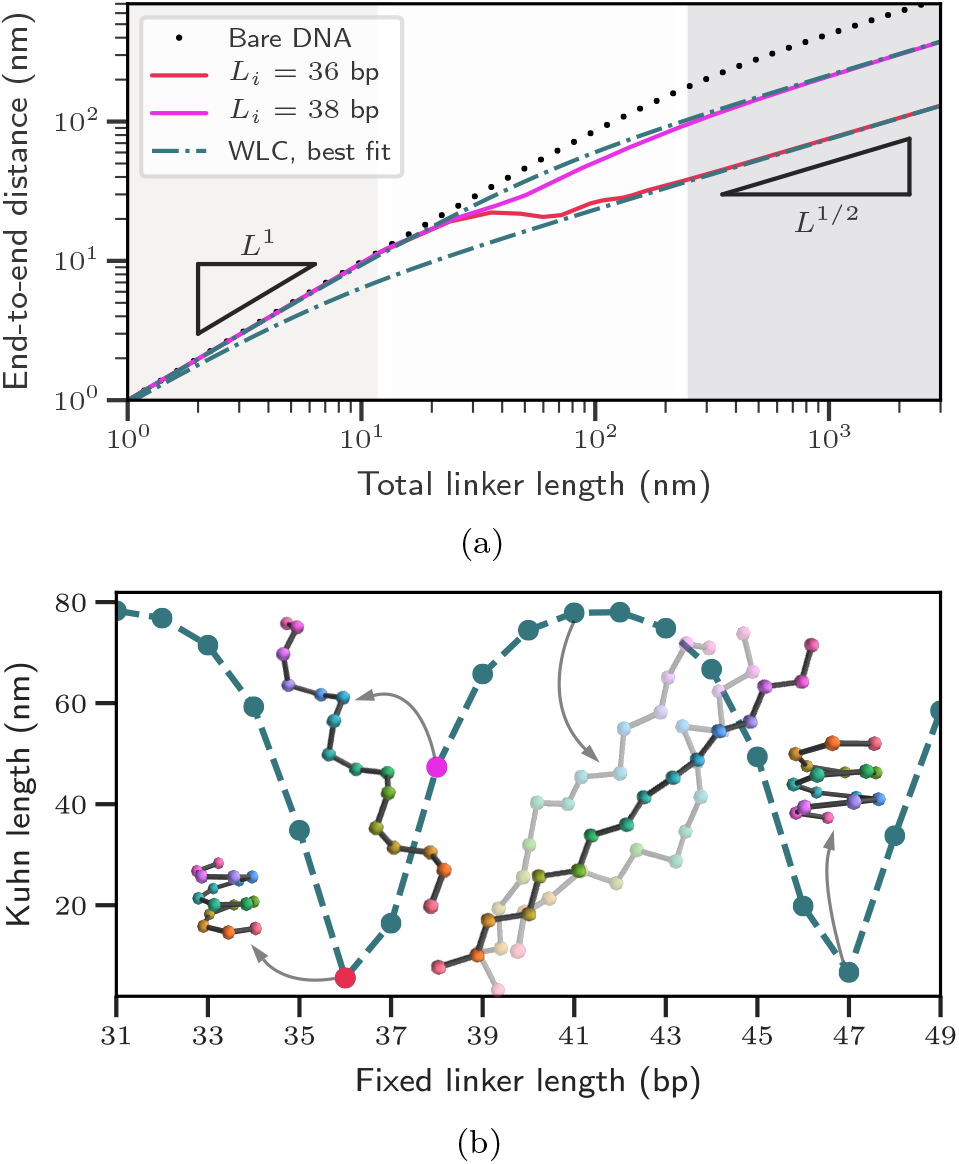
(a) Average end-to-end distances for homogeneous chromatin chains with 36 bp and 38 bp linkers, compared to bare DNA and the best-fit WLCs. Rigid rod, Gaussian chain, and cross-over regimes highlighted. (b) Kuhn lengths of homogeneous chromatin chains are 10.5bp periodic in linker length. Example minimum energy chain configurations are shown, including two example Monte Carlo (fluctuating) structures for the 41 bp case. Compact structures (36, 47 bp) afford more flexibility than less compact structures (41 bp).

To build a geometric intuition for how the kinks create this modified Kuhn length, we compare the Kuhn lengths of homogeneous, fluctuating chains to their zero-temperature configurations, where the entire chain is composed of rigid-rod linkers. Every homogeneous chain at zero temperature forms a helix of nucleosomes. The rise per basepair of the helix is determined by the spherical angles *θ* and *ϕ* connecting adjacent linkers (see Fig. 1). The nucleosome structure fixes *θ*, but *ϕ* depends linearly on the linker length, and is 10.5 bp periodic due to the DNA’s helicity. Select values of *ϕ* lead to more compact zero-temperature structures with a smaller rise per base-pair. As seen in Figure 2b, the corresponding fluctuating structures have smaller Kuhn lengths, and those with a larger rise per basepair at zero temperature have larger Kuhn lengths. The 10.5 bp periodicity of *ϕ* as linker length changes leads to the periodicity in Figure 2b. As *L*_*i*_ → ∞, the Kuhn length approaches that of bare DNA only slowly (see Supplemental Material [42], Fig. S4).

We next consider heterogeneous chains where the linker lengths are drawn uniformly from a range *μ* ± *σ*. In Fig. 3, we see that as we increase *σ*, the zero-temperature configuration of the chain interpolates between a helix at *σ* = 0 and a random walk at larger *σ*. As a result, the zero-temperature structure itself has a Kuhn length, which describes the compactness of the random walk. As in the homogeneous case, the Kuhn length of the zero-temperature chain qualitatively predicts that of the fluctuating structure, as seen in Fig. 3. We find that even a single basepair of variance in nucleosome positions (see e.g. Supplemental Material [42], Fig. S5) can create enough geometric stochasticity at zero-temperature to prevent the formation of regular fibers.

**FIG. 3.**
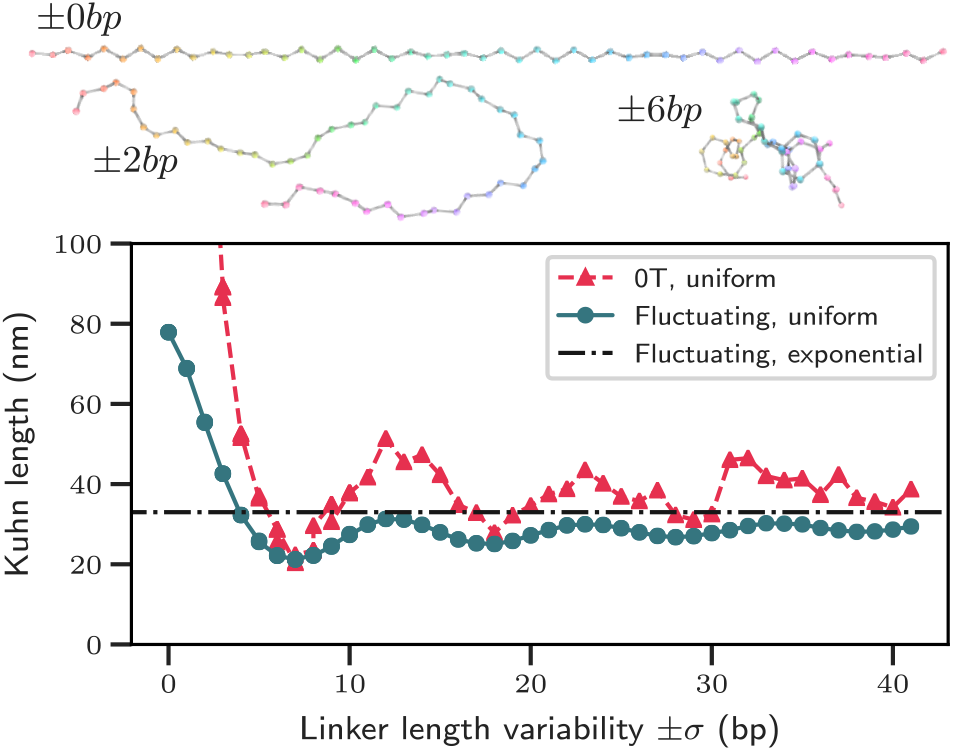
Kuhn length of a heterogeneous chromatin chain with uniformly distributed linker lengths chosen from the range *μ* ± *σ*, where *μ* = 41 bp. Example zero-temperature (0T) chains composed of rigid rod linkers are shown for *σ* = 0, 2, 6 bp. Kuhn length rapidly approaches that of the exponential chain (black line) in which linker lengths are exponentially distributed about the same *μ*.

In the cell, the simplest model of nucleosome binding is one in which nucleosomes bind uniformly randomly along the DNA [23]. In this model, the distances separating the nucleosomes are exponentially distributed (hereafter “the exponential chain”). While this picture ignores some details of *in vivo* nucleosome formation, Fig. 3 shows how any linker length distribution with sufficiently large variance (*σ*) will exhibit behavior similar to the exponential chain. Thus, the results that follow are likely robust to adding more detail to the nucleosome binding model.

When averaged over the distribution of possible nucleosome positions, the end-to-end distance of our exponential chain takes the form of a WLC with a rescaled Kuhn length (see Supplemental Material [42], Fig. S6). This is not wholly unsurprising, since at zero temperature our model differs from a freely rotating chain only in the correlation between linker length and *ϕ*, and the freely rotating chain is known to converge to a worm-like chain under appropriate scaling [50]. We extract the rescaled Kuhn length as a function of ⟨*L*_*i*_⟩ in Fig. 4.

**FIG. 4.**
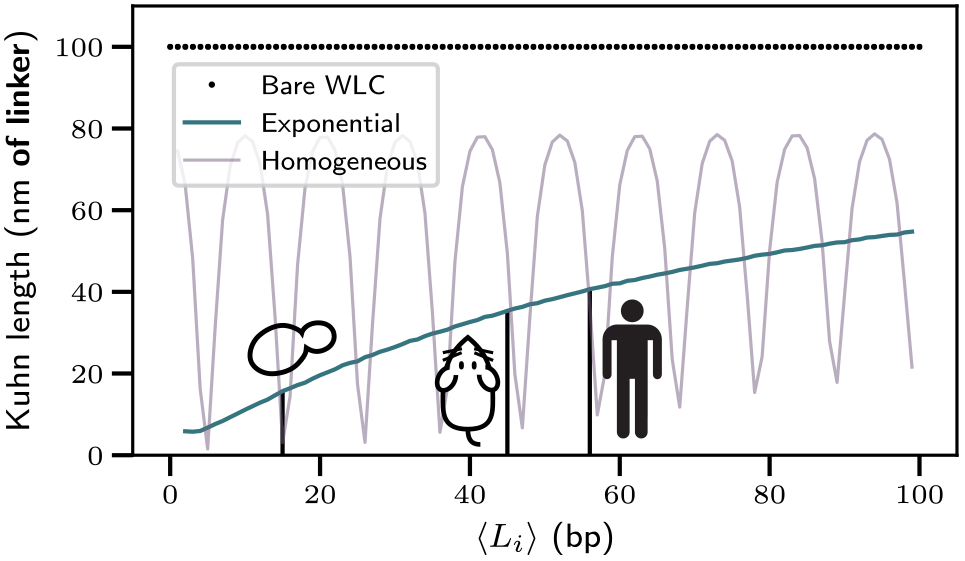
Kuhn lengths of chromatin chains with exponentially distributed linkers, as a function of the average linker length, which varies by cell type. Kuhn lengths for *S. cerevisiae* (⟨*L*_*i*_⟩ = 15 bp), mice embryonic stem cells (45 bp), and human T cells (56 bp) are marked.

Unlike the homogeneous case, where changing *L*_*i*_ selects between zero-temperature helices, increasing ⟨*L*_*i*_⟩ in the heterogeneous case scales the zero-temperature random walk. As a result, Fig. 4 lacks the 10.5 bp periodicity of the homogeneous chain. Thus, for the purposes of coarse-graining, an approximate knowledge of ⟨*L*_*i*_⟩ should be sufficient to capture chromatin’s average behavior as a WLC. A table of these Kuhn lengths is available in the Supplemental Material [42], Table S1.

While the Kuhn length describes the chain’s elasticity on long length scales, a more useful metric on shorter length scales is the probability of genomic contacts (*i.e.* the looping probability). By numerically inverting the Fourier transform in Eq. 3, we can analytically evaluate 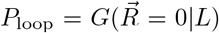, a modified *J*-factor with no orien-tational component. In Fig. 5, we plot this probability as a function of the loop size.

**FIG. 5.**
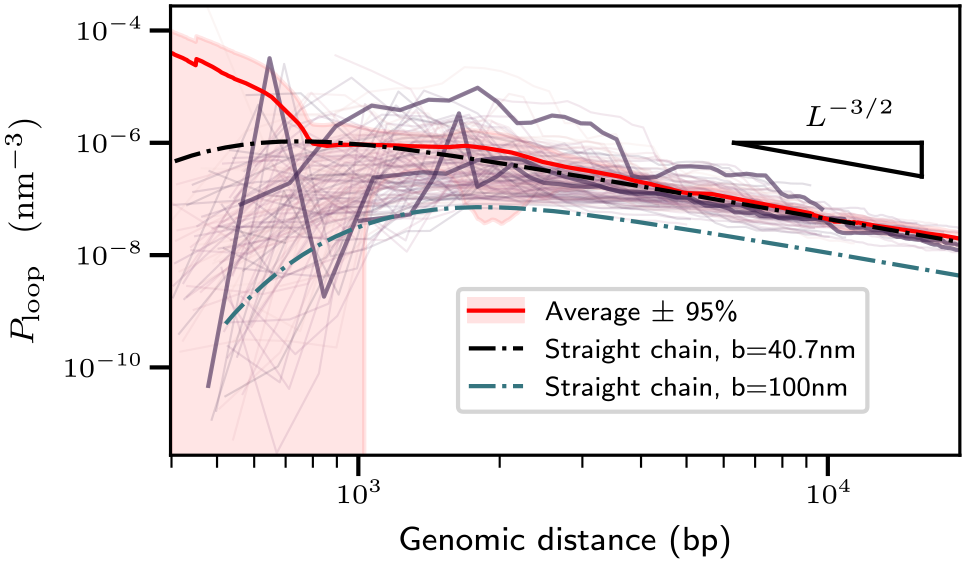
Looping probabilities as a function of genomic separation for exponential chains. Each purple line designates an individual chain with linker lengths drawn from an exponential distribution with *μ* = 56 bp. The red shaded area corresponds to 95% confidence intervals around the mean over the individual chains (red line). A wormlike chain (black dashed line) with the same Kuhn length as this average chain captures the looping probability for loops with at least three intervening nucleosomes. Three random individual chains are bolded.

Strikingly, we observe that for a fixed genomic separation, the difference between the most- and least-likely to loop chains spans up to six orders of magnitude depending on the intervening nucleosomes’ spacings. The average looping probability, which is relevant at timescales where nucleosome positions can re-randomize, is captured well by a single WLC. Due to this effective WLC’s reduced Kuhn length, we predict that chromatin’s propensity for forming sub-kilobase-scale loops should be one to four orders of magnitude larger than that of bare DNA. The predicted looping propensity peaks at a length scale typical of promoter contacts *in vivo* and consistent with Hi-C looping data [51] (see Supplemental Material [42], Fig. S8). This result highlights how even without models of DNA more detailed than the WLC— for example those including DNA melting [52–54]—the propensity of small DNA loops can be enhanced by proteins that promote kinks in the DNA if they are stochastically spaced.

Additionally, we find that the average looping probability is higher than that of most of the individual chains. Thus, even an “informationless” chromatin remodeler that merely promotes random nucleosome repositioning will greatly facilitate sub-kilobase loop formation. At longer length scales, the looping probabilities approach the characteristic *L*^−3/2^ Gaussian scaling. However, individual chains retain memory of their kinks, as indicated by how the highlighted chains persist above or below the average. Using other linker length distributions, such as uniformly distributed linkers, leads to qualitatively similar results (see Supplemental Materials [42], Fig. S9).

In conclusion, we provide rigorous justification for using an effective WLC to model *in vivo* chromatin. Due to the lack of experimental consensus on the persistence length *l*_*p*_ of chromatin [55], coarse-grained models of chromatin have historically used a range of values for *l*_*p*_, sometimes even just using that of bare DNA [56–58]. We show that this choice leads to at least a two-fold overestimation of the polymer’s stiffness and a several orders-of-magnitude underestimation of looping at short length scales. Some past models have extracted a parameter that describes the linear compaction of chromatin (bp per nm of fiber) from Hi-C looping probabilities (see [59, 60]). Our model’s parameter-free estimate of chromatin’s stiffness provides a theoretical explanation for the effective Kuhn length predicted from these experimental measurements.

Our model excludes various important facets of chromatin’s structure, such as interaction energies (sterics and stacking) and nucleosome “breathing”. Since nucleosome breathing simply corresponds to choosing a different distribution for the angle *θ* ∈ [0, *π*] between adjacent nucleosomes, incorporating breathing leaves our results qualitatively unchanged (see Supplemental Material [42], Fig. S10). However, a careful inclusion of breathing will likely require an explicit treatment of the effects of DNA sequence and linker histone on the nucleosome particle.

Our work highlights that the geometric effects of heterogeneous nucleosome binding dominate thermal fluctuations in determining chromatin’s elasticity *in vivo*. Our insights into chromatin’s stiffness will also inform future studies on the effects of loop extrusion factors [61] and epigenetic states [62] on chromatin organization.

Financial support for this work is provided by the National Science Foundation (NSF), Physics of Living Systems Program (PHY-1707751). Q.M. and B.B. acknowledge funding support from the NSF Graduate Fellowship program (DGE-1656518). B.B. acknowledges support from NIH Training Grant T32GM008294.

