## Supplemental Materials for "Heterogeneity in Nucleosome Spacing Governs Chromatin Elasticity"

July 19, 2019

Here, we provide a detailed derivation of the Green's function for a worm-like chain (WLC) with aperiodic defects, along with various checks to ensure the accuracy of our numerical schemes. First, we review some useful mathematical facts about rotations. We then provide a step-by-step derivation of the theory and a description of the numerical methods employed to perform the computations. For a more detailed description of the computations, all of our code (including code to generate all figures) can be downloaded from `https://github.com/brunobeltran/nuc_chain`.

In addition to supplementary figures referenced within the main text, we plot several internal and external controls designed to validate our numerical schemes.

**A Wigner D-function conventions** **7**

**B Supplementary Figures** **8**

*These authors contributed equally to this work.

### 1 The Kinked WLC

We model each linker in a chain of nucleosomes as a polymer strand whose conformation is given by the space curve $\vec{R}(s)$. The orientation of the polymer at each point along $\vec{R}(s)$ is represented by the orthonormal triad $\vec{t_i}(s)$ for $i = (1, 2, 3)$, where $\vec{t_3}$ is aligned with the tangent vector along the strand ($\vec{t_3} = \partial_s \vec{r}(s)$) and $\vec{t_1}$ and $\vec{t_2}$ are the material normals which live in the cotangent plane. We will denote this triad as $\Omega(s)$, which can also be interpreted as the rotation matrix from the lab frame to the frame defined by the material normals. Whenever we write this rotation element in terms of Euler angles, $\Omega(\alpha, \beta, \gamma)$ will correspond to the "z-y-z" Euler angles.

In order to represent the WLC's energy, we define the Euler vector $\vec{\omega} \in \mathfrak{so}(3)$ to be $\partial_s \vec{T} = \vec{\omega} \times \vec{T}$, where $T = \{\vec{t_1}, \vec{t_2}, \vec{t_3}\}$ is a vector with three elements. Because $\vec{t_3}$ is fixed tangent to the chain's path, $\omega_1(s)$ and $\omega_2(s)$ can be thought of as measuring the local bend in the chain, and $\omega_3(s)$ as measuring the twist at each point in the chain. In particular, notice that $\omega_1(s)^2 + \omega_2(s)^2 = \kappa(s)^2$, where $\kappa(s)$ is the curvature of $\vec{R}$.

#### 1.1 Defining the rigid kink

While the WLC's propagator fully describes the linker DNA in our model, we must also incorporate the effects of the nucleosome geometry on $\Omega(s)$. This is accomplished by treating the total orientational change from the point at which the DNA first binds the nucleosome to the last base pair bound as an instantaneous (discontinuous) change in $\Omega(s)$.

Loosely speaking, if we defined $\Omega_{\text{entry}}$ and $\Omega_{\text{exit}}$ to be the entry and exit orientations of DNA were we to explicitly model its structure as it wrapped around the nucleosome (as in Fig. 1), then we define $\Omega(s+\epsilon) = \Omega(s)\Omega_{\text{entry}}^{-1}\Omega_{\text{exit}}$.

To justify this formula, we recall how to compute the action of rotation matrices on coordinate frames. Suppose we have an orientation $\Omega_1(s)$ and a rotation $\Omega_2(\alpha, \beta, \gamma)$, both of which are represented as matrices in lab coordinates. In order to rotate $\Omega_1$ about the lab's x-, y- and z-axes with the Euler angles $\alpha$, $\beta$, $\gamma$, we multiply on the left to get the new orientation $\Omega_2\Omega_1$. In order to instead rotate

$$\Omega_1(s) = \begin{pmatrix} | & | & | \\ \vec{t_1}(s) & \vec{t_2}(s) & \vec{t_3}(s) \\ | & | & | \end{pmatrix}$$

about $\vec{t_1}(s)$, $\vec{t_2}(s)$ and $\vec{t_3}(s)$), we instead multiply on the right to get the new orientation $\Omega_1(s)\Omega_2$. Finally, in order to extract the rotation matrix that moves you between two known orientations, say from $\Omega_1(s)$ to $\Omega_2(s)$, in lab coordinates, we can use the usual change of coordinates formula $\Omega_{1\to 2} = \Omega_1^{-1}\Omega_2$.

Thus, we see that we are simply writing the effective entry to exit rotation in lab coordinates and multiplying from the right in order to impart the correct relative change in orientation to our chain.

Since the crystal structure of nucleosome-bound DNA has been solved, the orientations $\Omega_{\text{entry}}$ and $\Omega_{\text{exit}}$ can be determined experimentally. However, because the DNA forms a fairly uniform helix with an approximately constant intrinsic twist, it is equally easy to extract these orientations as a function of the number of base pairs bound.

In general, let $H(t) : [0, L] \to \mathbb{R}^3$ be the path traced out by the nucleosome-bound DNA. The intrinsic curvature $\kappa(t)$ and torsion $\tau(t)$ of the curve can be extracted from the Frenet-Serret frame, $\Omega_{\text{FS}} := \begin{pmatrix} \vec{u} & \vec{n} & \vec{b} \end{pmatrix}$ as

$$\frac{\partial}{\partial s}\begin{pmatrix} \vec{u} \\ \vec{n} \\ \vec{b} \end{pmatrix} = \begin{pmatrix} 0 & \tau(s) & -\kappa(s) \\ -\tau(s) & 0 & 0 \\ \kappa(s) & 0 & 0 \end{pmatrix}\begin{pmatrix} \vec{u} \\ \vec{n} \\ \vec{b} \end{pmatrix},$$

where $\vec{u}$, $\vec{n}$, and $\vec{b}$ are the tangent, normal, and binormal vectors to the curve. This means that in order to extract the final orientation of a material frame of reference $\Omega(s) = \begin{pmatrix} \vec{t_1} & \vec{t_2} & \vec{t_3} \end{pmatrix}$ that was transported with zero twist along the curve $H$, we need only compute the rotation matrix

$$H_{\text{rot}}(L) = \Omega_{\text{FS}}(0)^{-1}\Omega_{\text{FS}}(L)R_z(-\int_0^L \tau(s)ds), \tag{1}$$

where $R_z(\theta)$ is a rotation about the z-axis with angle $\theta$. In words, this says that we compute the effective entry and exit orientations of the Frenet frame, and then subtract off the total twist that we know the frame to have experienced by integrating the torsion.

For the case of the nucleosome, $H(t)$ is just a helix, which has constant torsion, so this simply reduces to $H_{\text{rot}}(L) = \Omega_{\text{FS}}(0)^{-1}\Omega_{\text{FS}}(L)R_z(L\tau_0 - L\tau))$ for an arbitrary amount of bound DNA $L$ and intrinsic bound DNA twist density $\tau_0$.

#### 1.2 Green's Function of TWLC

The molecular architecture of DNA results in a strand that opposes bending and twisting deformations. The simplest model that captures these effects is the (twistable) wormlike chain (TWLC) model. The elastic deformation energy of a WLC with twist,

$$\beta\mathcal{E} = \frac{l_p}{2}\int_0^L ds\ (\omega_1^2 + \omega_2^2) + \frac{l_t}{2}\int_0^L ds\ (\omega_3 - \tau)^2, \tag{2}$$

is quadratic in bend and twist. Here $l_p$ is the bend persistence length ($\approx 50\,\text{nm}$), $l_t$ is the twist persistence length ($\sim 100\,\text{nm}$), and $\tau$ is the natural twist density ($2\pi/(10.5\,\text{bp per turn of DNA})$). The polymer is also subject to the inextensibility constraint $|\partial_s \vec{r(s)}| = 1$ for all $s$.

The statistical behavior of the polymer strand is obtained by summing over all possible conformations and assigning each a Boltzmann weighting. Formally, the chain-orientation's Greens function is the conditional probability that a polymer of length $L$ will have fixed end orientation $\Omega(s = L)$ given that it has initial orientation $\Omega(s = 0)$, formally

$$G_0(\Omega|\Omega_0; L) = \int_{\Omega(s=0)}^{\Omega(s=L)} \mathscr{D}[\Omega(s)] \exp[-\beta\mathcal{E}] = \int_{\Omega_0}^{\Omega} \mathscr{D}[\Omega(s)] \exp[i \int_0^L ds\ \mathcal{L}(\Omega(s))]. \tag{3}$$

where the we can write the energy as an effective Lagrangian density $\mathcal{L} = il_p(\omega_1^2 + \omega_2^2)/2 + il_t(\omega_3 - \tau)^2/2$.

This path integral formulation implies a governing diffusion or "Schrödinger" equation of the form $\frac{\partial G_0}{\partial L} = \mathcal{H}_0 G_0$, where $\mathcal{H}_0 = \frac{1}{2l_p}\nabla^2_\Omega + \frac{1}{2}(\frac{1}{l_t} - \frac{1}{l_p})\frac{\partial^2}{\partial\gamma^2} + i\tau\frac{\partial}{\partial\gamma}$ and $\frac{\partial}{\partial\gamma}$ is the momentum operator in the third coordinate (conjugate to $\omega_3$). This equation has an explicit solution given by an eigenfunction expansion in Wigner D-functions $\mathcal{D}_l^{mj}(\Omega)$, namely

$$G_0(\Omega|\Omega_0; L) = \sum_{l=0}^{\infty} \sum_{m,j=-l}^{l} \mathcal{D}_l^{mj}(\Omega)\mathcal{D}_l^{mj*}(\Omega_0)\ \exp(-\lambda_l^j L), \tag{4}$$

where $\lambda_l^j = \frac{1}{2l_p}l(l+1) + \frac{1}{2}(\frac{1}{l_t} - \frac{1}{l_p})j^2 + i\tau j$. In other words, the Wigner functions satisfy $\mathcal{H}_0\mathcal{D}_l^{mj}(\Omega) = -\lambda_l^j\mathcal{D}_l^{mj}(\Omega)$. We now turn to the full end-to-end distribution function $G(\vec{R}, \Omega|\Omega_0; L)$, which gives the probability that a chain of length L that begins at the origin with fixed initial orientation $\Omega_0$ will end at position $\vec{R}$ with fixed end orientation $\Omega$. This amounts to restricting the solution in Eq. 3 to chains that end at position $\vec{R}$:

$$G_{\text{TWLC}}(\vec{R}, \Omega|\Omega_0; L) = \int_{\Omega_0}^{\Omega} \mathscr{D}[\Omega(s)] \exp[-\beta\mathcal{E}]\ \delta(\vec{R} - \int_0^L \vec{t_3} ds), \tag{5}$$

where $\delta$ is the Dirac delta. Upon Fourier transforming the position vector $\vec{R}$ to the wave vector $\vec{k}$ in Eq. 5, our problem becomes that of a TWLC with Hamiltonian $\beta\mathcal{H} = \beta(\mathcal{H}_0 + \mathcal{H}_{ext}) = \beta(\mathcal{H}_0 + i\vec{k}\cdot\vec{t_3})$. Since the original Hamiltonian is invariant to an arbitrary rotation in the lab frame, we choose $\vec{k}$ to point in the $\hat{z}$ direction, such that $\mathcal{H}_{ext} = k\hat{z}\cdot\vec{t_3} = k\cos\theta$. To obtain the full Green's function, we expand in the original basis of the Wigner functions

$$\hat{G}_{\text{TWLC}}(\vec{k}, \Omega|\Omega_0; L) = \sum_{lmj} \sum_{l_0 m_0 j_0} g_{l_0 m_0 j_0}^{lmj} \mathcal{D}_l^{mj}(\Omega)\mathcal{D}_{l_0}^{m_0 j_0 *}(\Omega_0), \tag{6}$$

To obtain the coefficients $g_{l_0 m_0 j_0}^{lmj}$, we utilize the property

$$\cos\theta\mathcal{D}_l^{mj} = \alpha_l^{mj}\mathcal{D}_{l-1}^{mj} + \beta_l^{mj}\mathcal{D}_l^{mj} + \alpha_{l+1}^{mj}\mathcal{D}_{l+1}^{mj}, \tag{7}$$

where $\alpha_l^{mj} = \sqrt{\frac{(l-m)(l+m)(l-j)(l+j)}{l^2(4l^2-1)}}$ and $\beta_l^{mj} = \frac{mj}{l(l+1)}$ are derived from the Clebsch-Gordon coefficients. Notice that $\cos\theta$ acts as a ladder operator which raises or lowers the $l$ index of the Wigner function; $m$ and $j$ are unperturbed, meaning $m = m_0$ and $j = j_0$. To simplify calculations, we consider the case where $m = 0$, which implies $\beta_l^{0j} = 0$. Plugging Eq. 6 into the new the "Schrödinger" equation $\frac{\partial \hat{G}}{\partial L} = \mathcal{H}_0 \hat{G} + ik \cos\theta \hat{G}$ amounts to solving the following tridiagonal ordinary differential equation:

$$\frac{\partial g_{l,l_0}^j}{\partial L} = -\lambda_l^j g_{l,l_0}^j + ik\alpha_l^j g_{l-1,l_0}^j + ik\alpha_{l+1}^j g_{l+1,l_0}^j \tag{8}$$

.

Solving Eq. 8 results in the Green's function in Fourier space for a TWLC, which models a single DNA linker in our nucleosome chain. In this work, we solve this ODE via matrix exponentiation after truncating $l_0 \in [0, 20]$ and $l \in [0, 50]$. A convergence analysis was conducted to ensure that increasing the maximum values of the indices $l_0$ or $l$ does not change the solution within a tolerance of $10^{-8}$. This approach matches our lab's previously published approach (see Figure 1.2), which relied on Laplace transforming equation 8 and expressing the solution in terms of continued partial fractions [1], but has the advantage of being significantly simpler to implement. This approach is called the "transfer matrix" formalism by the lab of Jie Yan, who applied a similar technique to single kinks previously [2].

#### 1.3 Green's Function of TWLC with Kinks

In order to incorporate the effects of the nucleosome rotation, we use the rotation matrix $\Omega_{\text{kink}} := H_{\text{rot}}$ derived in Section 1.1. Treating the rotation as a simple change of coordinates, we can rewrite Eq. 6 as

$$\hat{G}(\vec{k}, \Omega | \Omega_0; L) = \sum_l \sum_{l_0 j_0} g_{l\,l_0}^{j_0} \mathcal{D}_l^{0 j_0}(\Omega_{\text{kink}}^{-1} \Omega_f) \mathcal{D}_{l_0}^{0 j_0 *}(\Omega_0). \tag{9}$$

Next, we utilize the formula for the Wigner function of two successive rotations

$$\mathcal{D}_l^{0 j_0}(\Omega_{\text{kink}}^{-1} \Omega_f) = \sum_j \sqrt{\frac{8\pi}{2l+1}} \mathcal{D}_l^{0j}(\Omega_f) \mathcal{D}_l^{j j_0}(\Omega_{\text{kink}}^{-1}). \tag{10}$$

Renaming $j \to j_f$ and $l \to l_f$, the Green's function in Fourier space for a kinked TWLC becomes

$$\hat{G}(\vec{k}, \Omega_f | \Omega_0; L) = \sum_{l_f j_f} \sum_{l_0 j_0} B_{l_0, j_0}^{l_f, j_f} \mathcal{D}_{l_f}^{0 j_f}(\Omega_f) \mathcal{D}_{l_0}^{0 j_0 *}(\Omega_0), \tag{11}$$

where $B_{l_0,j_0}^{l_f,j_f} = \sqrt{\frac{8\pi^2}{2l_f+1}} \mathcal{D}_{l_f}^{j_f j_0}(-\gamma, -\beta, -\alpha) g_{l_f, l_0}^{j_0}$. Thus, each monomer in the nucleosome chain is uniquely defined by the length of the linker, $L$, and the rotation due to the nucleosome, $\Omega_{\text{kink}}$.

To compose multiple monomers in a chain, we utilize the composition property of Green's functions. In Fourier space, the spatial convolution of two

Green's function is just a product

$$G(\vec{k}, \Omega_2|\Omega_0; L_2 + L_1) = \int d\Omega_1 G(\vec{k}, \Omega_2|\Omega_1; L_2) G(\vec{k}, \Omega_1|\Omega_0; L_1) \tag{12}$$

This is functionally equivalent to multiplying the $B_{l_0,j_0}^{l_f,j_f}$ matrices in Fourier space.

#### 1.4 Kuhn Length Calculation

The Kuhn length is the single parameter that relates the average distance between two points on a chromosome to the amount of DNA that connects them. For a WLC, the kuhn length is $b = 2l_p$. For our kinked polymer, we utilize the standard definition $b = \lim_{n\to\infty} \langle R^2 \rangle / R_{max}$, where $R_{max}$ is the maximum length the polymer can be if all the linkers were perfectly straight (cumulative length of linker DNA).

The $m^{th}$ moment of the $z$ component of $R$ is given by

$$\lim_{k\to 0} \frac{\partial^m B_{00}^{00}}{\partial |k|^m} = (i)^m \langle R_z^m \rangle. \tag{13}$$

The only k-dependence in $B_{00}^{00}$ is in the linker propagator $g_{00}^0$, so to calculate $\langle R^2 \rangle$, we take derivatives with respect to k of Eq. 8. The resulting ODE can be efficiently computed in Laplace space, as has been described in [1].

To derive Eq. 13, we first notice that

$$\langle R \rangle = \langle e^{i\vec{k}\cdot \int_0^L ds\ \vec{t_3}} \rangle_{\Omega_0}^{\Omega_f} = \int d\Omega_0 \int d\Omega \sum_{lmj} \sum_{l_0 m_0 j_0} g_{l_0 m_0 j_0}^{lmj} \mathcal{D}_l^{mj}(\Omega) \mathcal{D}_{l_0}^{m_0 j_0 *}(\Omega_0) = g_{00}^{00}. \tag{14}$$

The last equality comes from integrating over the start and end orientations. Due to the orthogonality property of Wigner functions, this selects out the $g_{00}^0$ coefficient. From Eq. 14, we can compute higher moments of $R$ by noticing that $\frac{\partial^m g_{00}^0}{\partial |k|^m} = \langle i^m R_z^m exp(ikR_z) \rangle$ since $\vec{k} = k\hat{z}$, which produces the result in Eq. 13.

#### 1.5 Looping Probability Calculation

To compute the real space green's function, we compute the inverse Fourier transform of Eq. 11:

$$\begin{aligned} G(\vec{R}, \Omega|\Omega_0; L) &= \frac{1}{(2\pi)^3} \int_0^\infty d\vec{k}\ \hat{G}(\vec{k}, \Omega|\Omega_0; L) \exp(-i\vec{k} \cdot \vec{R}) \\ G(\vec{R}; L) &= \frac{1}{(2\pi)^3} \int_0^\infty d\vec{k}\ G(\vec{k}; L) \exp(-i\vec{k} \cdot \vec{R}) \\ &= \frac{1}{2\pi^2} \int_0^\infty dk\ k^2 j_0(kR) B_{00}^{00}(k; L) \end{aligned} \tag{15}$$

where $j_0(kR)$ is the $0^{th}$ spherical bessel function. The second equality comes from integrating over all possible fixed end orientations. To perform the Fourier inversion, we chose 30,000 values of $K = k/(2l_p)$ between 0 and $10^5$, in which 20,000 are linearly spaced and 10,000 are log-spaced. The integration was performed using cubic spline interpolation between these points. We elected not to use standard adaptive quadrature schemes because those available via the scipy package did not converge in a reasonable amount of time.

To determine the bounds on $K$, we examined the numerics of the propagator for short linkers less than 15 bp. Although $K = 10^5$ is not high enough in this limit, we find that higher $K$ values only contribute minimally to the overall value of the integral. At high $K$ and low $N = L/(2l_p)$, the periodicity of the integrand is dictated by the sinc function $\sin(KN)/(KN)$. Thus, we chose there to be 20,000 linearly spaced points such that there would be at least 12 $k$ values per period for a homogeneous chain with 50 bp linkers. For low $K$ and high $N$, the integrand has a shorter period than the sinc function; however, the propagator for longer linkers rapidly falls off to zero, so the 10,000 log-spaced $K$ points sufficiently capture this limit.

The probability that two ends of the nucleosome chain come into contact is obtained by evaluating the Green's function at $R = 0$: $P_{\text{loop}} = G(\vec{R} = 0; L)$, where $P_{\text{loop}}$ is in units of $L^{-3}$. We recognize that this looping "concentration" is equivalent to a $J$ factor, ignoring end orientations. At long genomic distances, the looping probability of our kinked polymer matches that of a Gaussian chain with $P(R = 0; N) = (\frac{3}{2\pi N b^2})^{3/2}$, where $b$ is the Kuhn length and $N$ is the number of Kuhn lengths. Note, this corresponds to a power law scaling of $N^{-3/2}$. Renormalizing such that length is expressed in units of basepairs instead of Kuhn lengths, the Gaussian looping probability becomes $P(R = 0; R_{max}) = (\frac{3}{2\pi R_{max} b})^{3/2}$.

### A Wigner D-function conventions

Throughout our derivations, we define our Wigner functions $\mathcal{D}_l^{mj}$ to contain the following scaling factor

$$\mathcal{D}_l^{mj} = \sqrt{\frac{2l+1}{8\pi^2}} D_l^{mj}$$

Here, $\mathcal{D}_l^{mj}$ refers to our renormalized Wigner D's, and $D_l^{mj}$ refers to the conventional wigner D functions. This renormalization results in the following orthonormality condition:

$$\int_0^{2\pi}\int_0^{2\pi}\int_0^{\pi} \mathcal{D}_l^{mj*}(\alpha,\beta,\gamma)\mathcal{D}_{l_0}^{m_0 j_0}(\alpha,\beta,\gamma) d\alpha \sin\beta d\beta d\gamma = \delta_{m,m_0}\delta_{j,j_0}\delta_{l,l_0}$$

Thus we can reinterpret our redefined Wigner D functions are unnormalized distribution functions:

$$\int_0^{2\pi}\int_0^{2\pi}\int_0^{\pi}\mathcal{D}_0^{00}(\alpha,\beta,\gamma)d\alpha\sin\beta d\beta d\gamma = \sqrt{8\pi^2}.$$

This scaling factor is chosen so that the D-functions satisfy the following ladder operation condition:

$$\cos\theta\mathcal{D}_l^{mj} = \alpha_l^{mj}\mathcal{D}_{l-1}^{mj} + \beta_l^{mj}\mathcal{D}_l^{mj} + \alpha_{l+1}^{mj}\mathcal{D}_{l+1}^{mj},$$

where $\alpha_l^{mj} = \sqrt{\frac{(l-m)(l+m)(l-j)(l+j)}{l^2(4l^2-1)}}$ and $\beta_l^{mj} = \frac{mj}{l(l+1)}$ are derived from the Clebsch-Gordon coefficients.

To combine two successive Wigner D rotations together ($\Omega_1$ followed by $\Omega_2$),

$$\sqrt{\frac{2l+1}{8\pi^2}}\mathcal{D}_l^{mj}(\Omega_2\Omega_1) = \sum_\mu \mathcal{D}_l^{m\mu}(\Omega_1)\mathcal{D}_l^{\mu j}(\Omega_2)$$

The connection between our Wigner D's and Spherical Harmonics is as follows ($j = 0$):

$$\sqrt{2\pi}\mathcal{D}_l^{m0}(\alpha,\beta,\gamma) = Y_l^{m*}(\beta,\alpha)$$

### B Supplementary Figures

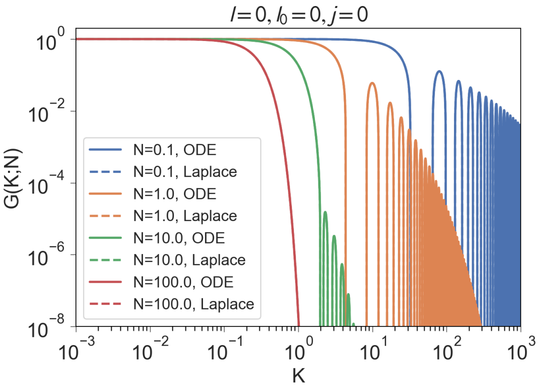

(a)

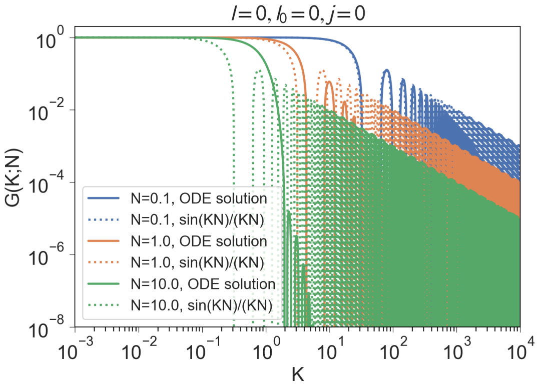

(b)

Figure 1: Coefficient of the TWLC linker propagator in Fourier space $g_{00}^{0}(K;N)$ as a function of the non-dimensionalized variable $K = k/(2l_p)$, where $N = L/(2l_p)$ is the chain length in number of Kuhn lengths. (a) The solution to the ODE using Pade approximation ('ODE') matches the solution obtained via partial fractions ('Laplace') in [1]. (b) Comparison of ODE solution to the propagator for a rigid rod, which is given by $\sin(KN)/(KN)$. The solutions agree for small $N$, as expected.

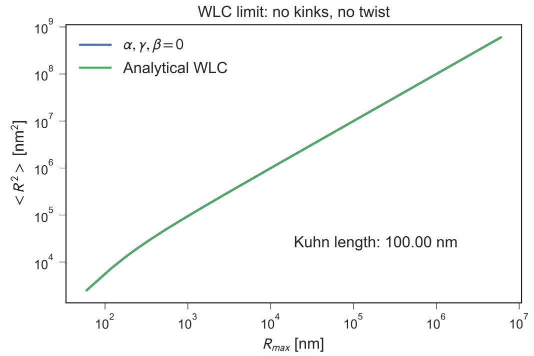

(a)

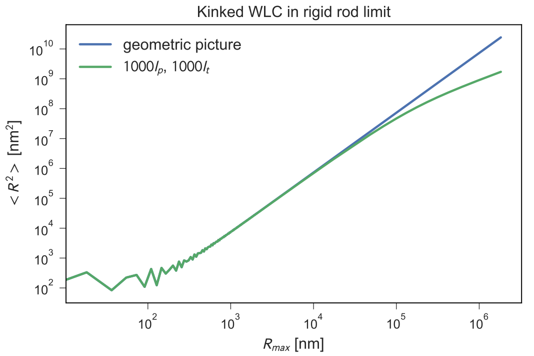

(b)

Figure 2: (a) When the Euler angles are set to zero, the nucleosome chain is equivalent to a bare WLC with no kinks. We demonstrate the kinked theory matches the analytical formula for a WLC: $\langle R^2 \rangle = 2 l_p R_{max} [1 - \frac{l_p}{R_{max}} (1 - e^{-R_{max}/l_p})]$. (b) As $l_p \to \infty$, each DNA linker becomes a rigid rod, in which case $\langle R^2 \rangle = \langle R_{max}^2 \rangle$. We demonstrate that the theory matches the purely geometric picture in this limit.

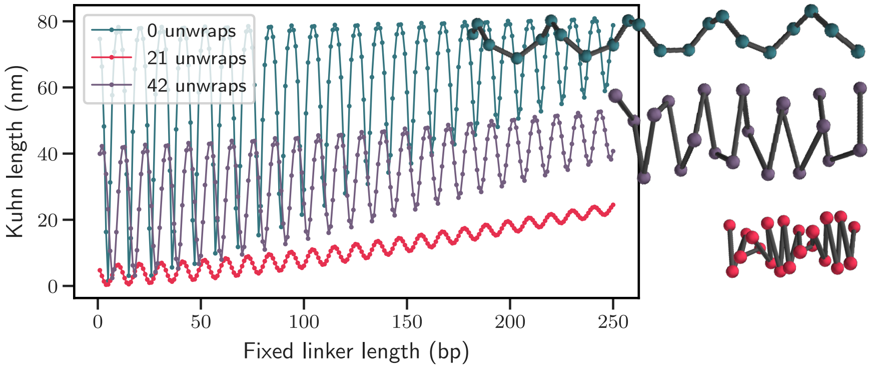

Figure 3: Kuhn lengths of homogeneous chains as a function of the fixed linker length for different unwrapping levels. In the main text, we assume all 147 bp of DNA are bound to the histone octamer. However, some nucleotides may "unwrap" from the octamer due to nucleosome breathing or active remodeling. Because the octamer must align with the major groove of the DNA double helix, the most common unwrapping amounts are multiples of 10.5 bp from the entry and exit sides. Here, we plot two common unwrapping levels — 10.5 bp on each side and 21 bp on each side. We find that the 10.5 bp periodicity in the Kuhn length is maintained at different wrapping levels, but the quantitative values change because the kink angle $\theta$ connecting adjacent nucleosomes changes.

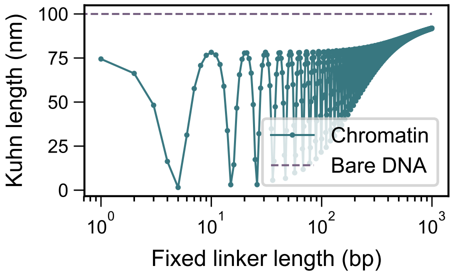

Figure 4: Kuhn lengths of homogeneous chains in the limit of long linker lengths. The Kuhn length obeys the same power law scaling as the kink density, which goes as $1/L$ where $L$ is the linker length. In this limit, fluctuations overpower the geometrical effects of nucleosomes.

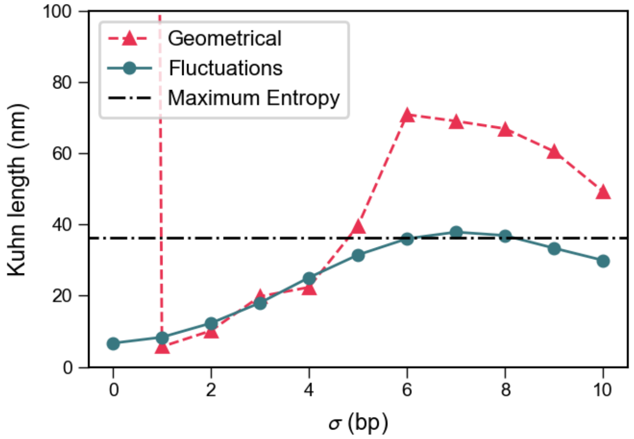

Figure 5: Kuhn length of a heterogeneous chromatin chain with uniformly distributed linker lengths chosen from the range $\mu \pm \sigma$, where $\mu = 47\,\mathrm{bp}$. For this choice of $\langle L_i \rangle$, the Kuhn length of the "geometrical" chain (composed of rigid rods) matches that of the fluctuating chain with only $\sigma = 1\,\mathrm{bp}$ of variability. With increasing $\sigma$, both the geometrical and fluctuating chains rapidly converge to the Kuhn of the maximum entropy chain, in which linker lengths are exponentially distributed about the same $\mu$.

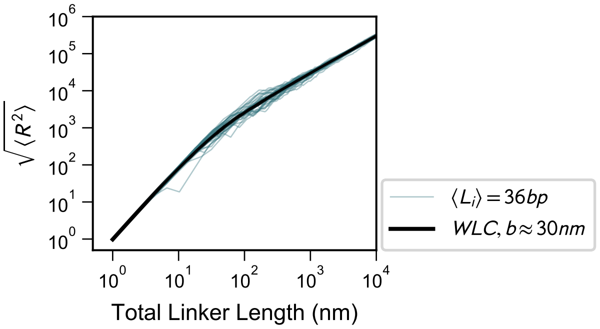

Figure 6: Average end-to-end distances of many chromatin chains with exponentially distributed linker lengths, as compared to the best-fit wormlike chain. Unlike the homogeneous chain case, both the short and large size scales are well explained by a single effective TWLC.

| $\langle L_i \rangle$ | $b_{\mathbf{linkers}}$**(nm)** | $\langle L_i \rangle$ | $b_{\mathbf{linkers}}$**(nm)** | $\langle L_i \rangle$ | $b_{\mathbf{linkers}}$**(nm)** |
|---|---|---|---|---|---|
| 10 | 11.14 | 35 | 29.78 | 60 | 42.08 |
| 11 | 12.00 | 36 | 30.24 | 61 | 42.70 |
| 12 | 12.87 | 37 | 30.74 | 62 | 42.90 |
| 13 | 13.62 | 38 | 31.52 | 63 | 43.38 |
| 14 | 14.68 | 39 | 32.06 | 64 | 43.75 |
| 15 | 15.69 | 40 | 32.62 | 65 | 44.20 |
| 16 | 16.40 | 41 | 33.09 | 66 | 44.51 |
| 17 | 17.17 | 42 | 33.87 | 67 | 44.88 |
| 18 | 17.76 | 43 | 34.16 | 68 | 45.36 |
| 19 | 18.75 | 44 | 34.74 | 69 | 45.66 |
| 20 | 19.59 | 45 | 35.36 | 70 | 46.00 |
| 21 | 20.30 | 46 | 35.91 | 71 | 46.40 |
| 22 | 21.04 | 47 | 36.32 | 72 | 46.85 |
| 23 | 22.01 | 48 | 36.95 | 73 | 47.00 |
| 24 | 22.61 | 49 | 37.22 | 74 | 47.42 |
| 25 | 22.99 | 50 | 37.84 | 75 | 47.68 |
| 26 | 24.00 | 51 | 38.21 | 76 | 47.94 |
| 27 | 24.55 | 52 | 38.69 | 77 | 48.32 |
| 28 | 25.33 | 53 | 39.19 | 78 | 48.84 |
| 29 | 25.86 | 54 | 39.64 | 79 | 49.11 |
| 30 | 26.50 | 55 | 40.10 | 80 | 49.30 |
| 31 | 27.20 | 56 | 40.66 | 81 | 49.68 |
| 32 | 27.99 | 57 | 41.01 | 82 | 50.13 |
| 33 | 28.34 | 58 | 41.39 | 83 | 50.24 |
| 34 | 29.16 | 59 | 41.98 | 84 | 50.50 |

Table 1: We report the Kuhn lengths of heterogenous chromatin chains with exponentially distributed linker lengths as a function of the average linker length $\langle L_i \rangle$. $b_{\mathbf{linkers}}$ is in units of cumulative linker length. In order to correctly simulate a chromatin chain where nucleosomes are free to re-position themselves, first simulate a WLC (or simply a Rouse chain for long length scales) with the Kuhn length provided. In order to convert distance along the chain back into units of genomic distance, use the formula $L_{\text{genomic}} = L_{\text{linkers}}(1 + 146/\langle L_i \rangle)$.

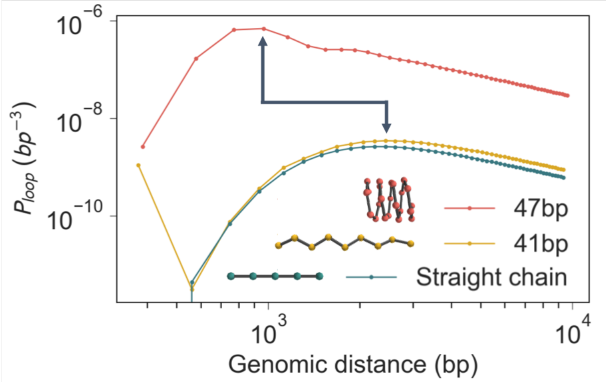

Figure 7: Looping probability as a function of genomic distance, starting from 2 nucleosomes down the chain (numerics are unstable for the first nucleosome). For linker lengths with compact, zero-temperature configurations (47 bp), the peak in the looping probability occurs at around a kilobase, representing a shift to the left compared to a perfectly straight chain of nucleosomes connected by bare WLCs with Euler angles set to zero. This is due to short range contacts between nucleosomes on adjacent rungs of the chromatin superhelix. Finally, notice that in log-log space, a vertical shift in the looping probability corresponds to a smaller effective Kuhn length in the long chain limit.

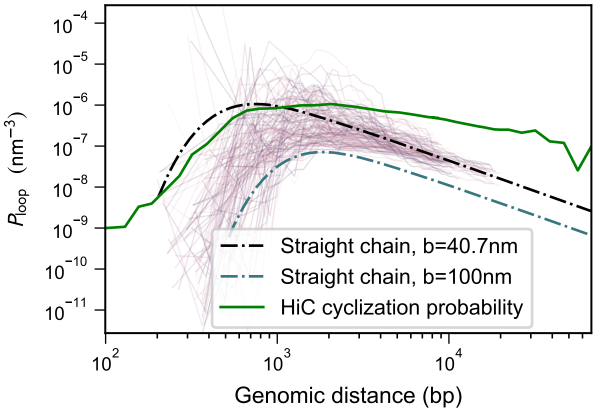

Figure 8: A simplified version of Fig. 5 from the main text. Each purple line designates the looping probability as a function of genomic distance of an exponential chain with $\mu = 56\,\text{bp}$. The dotted black line is the best-fit WLC with Kuhn length predicted by our $\langle R^2 \rangle$ calculation. The green curve is the Hi-C cyclization probability (Fig. 1 in [3], NcoI restriction). The green curve (originally in read-count units) has been rescaled to match the peak of the black curve. The experimental looping probability first reaches its maximum at the sub-kilobase length scale predicted by our model. While our theoretical looping curve immediately begins to fall off due to the Rouse behavior of the chain at longer lengths scales, the cyclization data remains relatively flat, and eventually begins to fall off at a much slower rate. This deviation from our theoretical predictions at longer length scales is expected due to interactions between distal segments of chromatin that are not included in our model. Future work that incorporates supercoiling, loop extrusion factors, and epigenetic interactions, and other long-scale chromatin interactions, can use a wormlike chain with our predicted persistence length to correctly capture this longer-scale looping behavior.

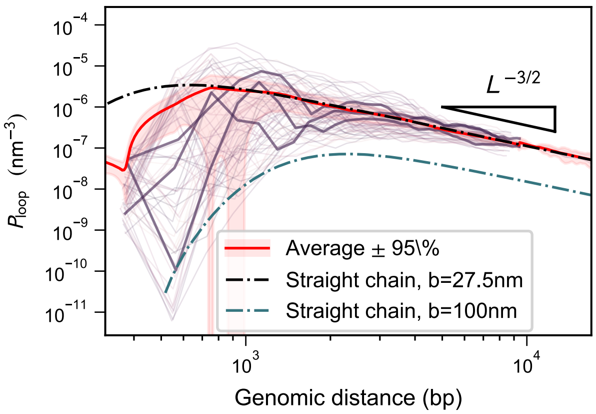

Figure 9: Looping probability as a function of genomic separation for chains with uniformly distributed linkers between 31 and 52 bp. The red shaded area corresponds to 95% confidence intervals around the mean over the individual chains (red line). A WLC with Kuhn length predicted by our $\langle R^2 \rangle$ calculation (black dotted line) still captures this average looping probability for chains with at least 3 intervening nucleosomes.

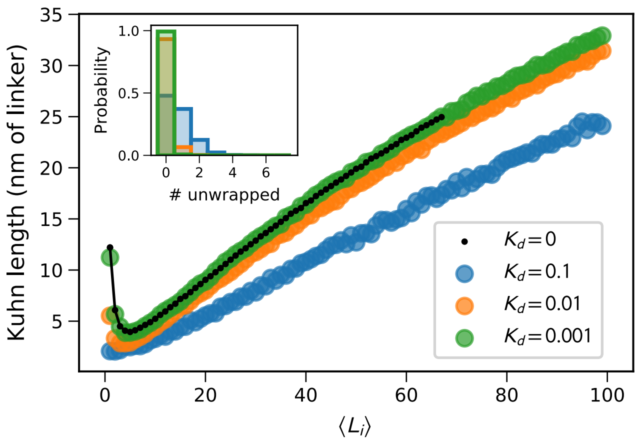

Figure 10: The Kuhn length of exponential chains as a function of mean linker length is plotted (as in Figure 4) for a toy model where the major groove of the DNA binds the histone octamer at 14 sites spaced 10.5 base pairs apart, and all 14 sites have fixed binding and unbinding rates $k_{\text{on}}$ and $k_{\text{off}}$, respectively. The three example curves represent three log-spaced values of $K_d = k_{\text{off}}/(k_{\text{on}} + k_{\text{off}})$, which corresponds to linearly-spaced changes in the dissociation energy of each contact site. The inset shows the distribution of nucleosomes that have 0, 1, 2, or more sites unbound for each value of $K_d$ plotted. The qualitative trend remains the same, and adding non-trivial unwrapping merely increases the flexibility of the nucleosome chain even further.
